# ChAHP Silences SINE Retrotransposons by Inhibiting TFIIIB Recruitment

**DOI:** 10.1101/2025.07.02.662776

**Authors:** Jakob Schnabl-Baumgartner, Fabio Mohn, Michaela Schwaiger, Josip Ahel, Jennifer Steiner, Yukiko Shimada, Sirisha Aluri, Marc Bühler

## Abstract

Short interspersed nuclear elements (SINEs) are abundant non-autonomous transposable elements derived from RNA polymerase III (POL III)-transcribed short non-coding RNAs. SINEs retain sequence features recognized by the POL III machinery and constitute a substantial portion of vertebrate genomes. Despite their impact on genome stability and evolution, the mechanisms governing SINE transcription remain poorly understood. Although DNA methylation and heterochromatin formation have been implicated in their repression, we find these pathways play only a minor role in mouse embryonic stem cells. Instead, we identify the ChAHP complex as a key repressor of SINE B2 elements. ChAHP directly inhibits POL III transcription by blocking TFIIIB recruitment without affecting TFIIIC binding. This selective interference prevents transcription initiation and highlights a distinct regulatory mechanism. Our findings establish ChAHP as a non-canonical repressor of POL III-dependent SINE transcription, offering new insights into the control of this pervasive class of non-coding genomic elements.

## INTRODUCTION

Transposable elements (TEs) are mobile genetic elements present in nearly all eukaryotic genomes. These “selfish” DNA sequences propagate by hijacking the cellular machinery of their hosts, enabling their mobilization and replication within genomes. Retrotransposons, a major subclass of TEs, employ an RNA intermediate and reverse transcriptase (RT) to increase their copy number via a copy-and-paste mechanism ^1–3^. These copies constitute substantial portions of vertebrate genomes; for instance, retrotransposons account for approximately 37% of the mouse genome ^4^.

TE activity can have profound implications for host genomes. Transcription of TEs can alter expression of nearby genes, disrupt gene-regulatory networks and potentially lead to developmental defects and diseases. Further, TEs can serve as hotspots for recombination and chromosomal rearrangements, and when mobilized, can disrupt gene function by insertional mutagenesis, both contributing to genomic instability. In contrast to negatively impacting their host’s fitness, TEs have also been associated with phenotypic variation, environmental adaptation, and the acquisition of new regulatory sequences. TEs are therefore considered important drivers of genome evolution ^3,5–7^.

To mitigate the deleterious effects of TEs, hosts have evolved diverse mechanisms to repress their activity. These include heterochromatin formation, DNA methylation, or small RNA-mediated silencing. The repressive activities are typically directed to TEs via sequence-specific recognition mechanisms and are mediated by multi-protein complexes such as KRAB-ZNFs, HUSH, or piRISC ^8–10^. These repressive mechanisms have been studied largely in the context of several kb long, RNA Polymerase II (Pol II) transcribed autonomous TEs such as endogenous retroviruses (ERVs) or long interspersed nuclear elements (LINEs). In contrast, the regulation of the highly abundant class of RNA Polymerase III (POL III)-transcribed, non-autonomous short interspersed nuclear elements (SINEs) is much less understood.

SINEs are chimeric elements with homology to endogenous short non-coding RNAs and LINE elements, present with over one million insertions in the mouse genome. They originate from cellular RNAs such as tRNAs (e.g., the mouse SINE B2 family), 7SL RNAs (e.g., mouse SINE B1/Alu) or 5S rRNAs (e.g., SINE3 in zebrafish)^11–13^. As a result, SINEs are relatively short, typically ranging from 80-300 bp. In addition to the 5ʹ region, which contains POL III promoter elements, active SINEs contain a genomically encoded poly(A)-stretch and often a short 3ʹ region that is partially homologous to the 3ʹ untranslated region (UTR) of LINEs. SINEs do not encode proteins; therefore, their propagation within the host genome depends on the RT and integration machinery encoded in LINEs, which recognize the 3ʹ end of SINE transcripts ^14,15^.

SINEs in the mouse genome can be classified in five families, namely B1/Alu, B2, B4/RSINE, ID and MIR (Figure S1A). They all contain so-called type 2 POL III promoters, like the highly abundant Alu and MIR families in humans. These internal promoters consist of A- and B-box elements that are directly recognized by the transcription factor complex TFIIIC, composed of six subunits (GTF3C1-GTF3C6) ^16,17^. TFIIIC binding leads to subsequent recruitment of TFIIIB, comprising TBP, BRF1, and BDP1, which in turn enables the binding of POL III and the initiation of transcription^18^. Unlike Pol II-mediated transcription, which involves multiple regulatory steps from recruitment to pre-initiation complex (PIC) formation and productive elongation, POL III transcription is believed to start immediately following the successful recruitment of POL III PIC ^19–21^. However, in contrast to the well-characterized canonical POL III-transcribed genes, transcriptional regulation at SINEs remains poorly understood. Early studies on SINEs have detected SINE transcripts in multiple different mouse tissues as well as during embryonic development ^22–24^. More recent studies in mouse and human cells revealed that multiple SINE insertions are bound by the POL III machinery, indicating that they are actively transcribed ^25–28^. Moreover, evolutionary young B1 and B2 SINEs show high numbers of polymorphic insertions when comparing different mouse strains, indicating that these SINEs have retained the ability to retrotranspose ^29^. Hence, host organisms must have evolved means to restrict further spreading. Both DNA methylation and heterochromatin formation have been implicated in SINE transcriptional silencing ^30–32^, yet the precise role of these repressive pathways in SINE regulation has remained obscure. In addition, recent work revealed that SINE elements are upregulated upon perturbation of the chromatin regulatory ChAHP complex consisting of the DNA binding factor ADNP, chromatin remodeler CHD4 and heterochromatin binding HP1 proteins ^33,34^. Like for the other repressive pathways, it remained unknow whether this is an indirect effect or a direct consequence of ChAHP impeding POL III transcription at SINE insertions.

We therefore set out to elucidate if and how SINE elements are regulated at the transcriptional level. We demonstrate that DNA and H3K9 methylation play only a minor role in repressing SINEs in mouse embryonic stem cells. Instead, we find that ChAHP selectively interferes with the recruitment of TFIIIB to the promoter of SINE B2 elements, thereby preventing initiation of transcription by POL III.

## RESULTS

### Evolutionary young SINEs are actively transcribed by POL III in mESCs

Given the ∼1.4 million SINE insertions in the mouse genome, quantifying expression of individual SINEs by RNA-seq is challenging because of the high sequence similarity of individual copies. Furthermore, the vast majority of reads mapping to SINE sequences originate from intronic insertions within expressed protein-coding genes that are transcribed by Pol II, making it difficult to correlate changes of RNA levels with changes in POL III transcription ^4^ (Figure S1B). Therefore, we first defined genomic loci transcribed by POL III using ChIP-seq for the core subunit POLR3A in mouse embryonic stem cells (mESC). Besides the expected POL III targets such as tRNA, U6 RNA, or 5S rRNA genes, most POL III peaks (n=2400, >80% of total peaks) locate over SINE insertions (Figure 1A-C). Consistent with previous observations, the POL III-bound SINE copies are strongly enriched for evolutionary young elements containing intact A- and B-boxes ^26^ (Figure S1C). Notably, these putatively active SINEs belong to the B1_Mm, B1_Mus1, B1_Mus2, and B2_Mm1a, B2_Mm1t, B2_Mm2 subfamilies which can actively retrotranspose in mice ^29^ (Figure 1B, S1D). Due to their high copy number in the genome and the likely non-random insertion mechanism, SINEs are often directly adjacent or even nested with other repeat insertions, complicating the unambiguous assignment of POL III peaks to individual elements ^35,36^ (Figure S1E). To avoid mis-assignments of ChIP peaks, we excluded all SINE insertions within 200 bp of other SINEs or POL III transcription units. Despite reducing the number of analyzable loci, this refined strategy identified a similar proportion of SINE-associated peaks, reinforcing the specificity of the high enrichment of SINEs (Figures 1C, S1D, S1F). For stringency and consistency, we have used this unambiguous SINE set for further analysis.

**Figure 1.**
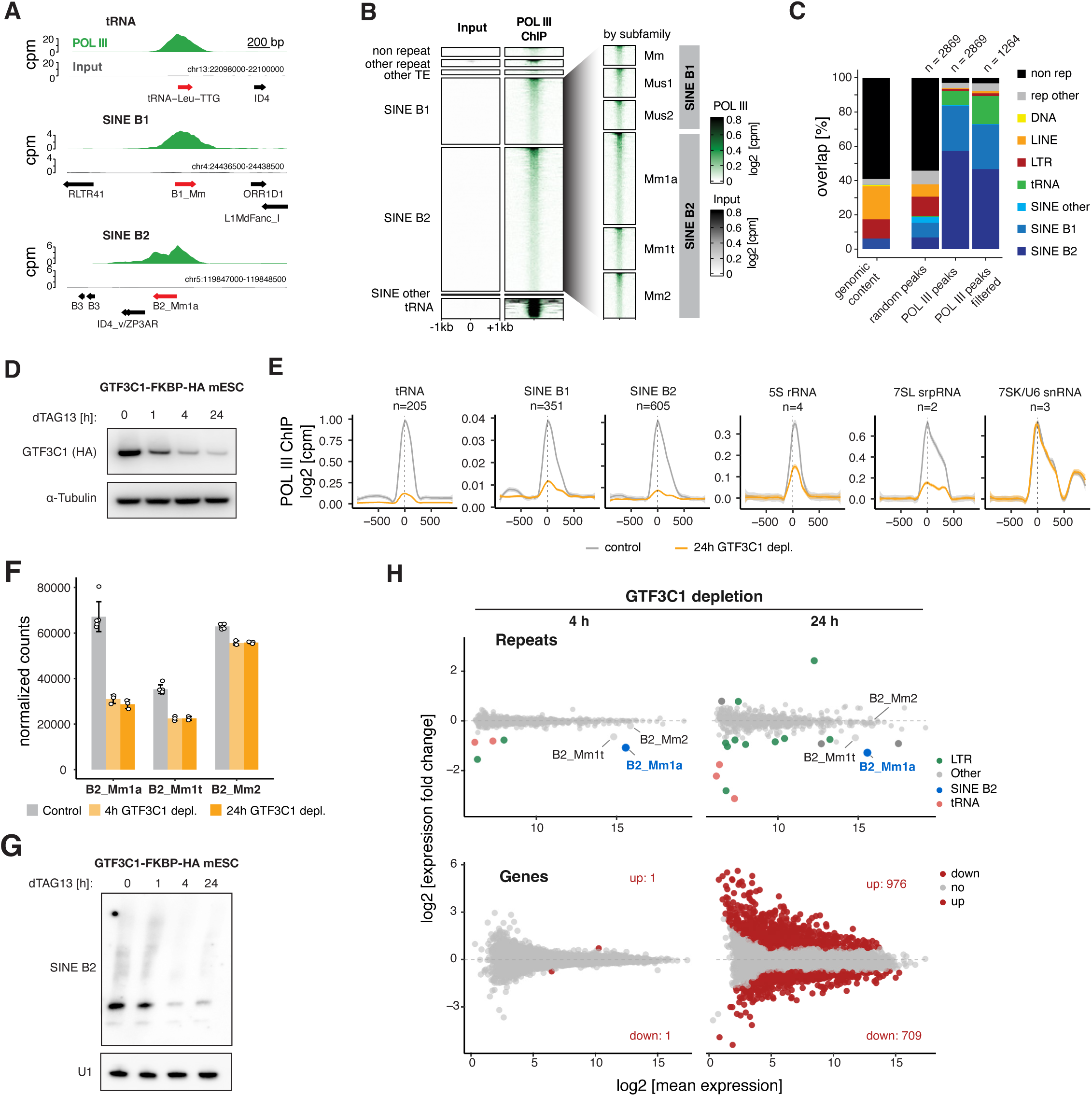
Evolutionary young SINEs are actively transcribed by POL III in mESCs. **(A)** Genome browser tracks showing POL III ChIP-seq signal at representative genomic loci: tRNA (top), SINE B1 (middle), and SINE B2 (bottom). Input signal shown in gray, POL III signal in green. Repeat annotations below each track. **(B)** Heatmaps of POL III ChIP-seq enrichment at peaks split by repeat family (left) or SINE B1 and SINE B2 subfamilies (right). **(C)** Overlap of POL III ChIP-seq peaks (right) with genomic annotations compared to the overall genomic content of these features (left). **(D)** Immunoblot of GTF3C1-FKBP-2xHA protein depletion following 500nM dTAG13 treatment for the indicated times. α-Tubulin serves as loading control. **(E)** Meta-profiles of POL III ChIP-seq signal (log2[cpm]) across different RNA Polymerase III-transcribed elements that overlap a POL III peak, in control (gray) and 24h GTF3C1-depleted (yellow) conditions. The number of elements (n) used to generate meta-profiles is indicated. **(F)** Quantification of normalized SINE-B2 subfamily transcript levels in control and GTF3C1 depletion conditions at indicated time points. Data represent mean ± SD from three independent replicates. **(G)** Northern blot analysis of SINE B2 RNA following GTF3C1 depletion for the indicated times. U1 snRNA serves as loading control. **(H)** MA plots showing differential expression of retrotransposons (upper panels) and genes (lower panels) after 4 h (left) and 24 h (right) of GTF3C1 depletion. Significantly regulated genes and repeats are highlighted in color, SINE B2 subfamilies are labeled.

To determine whether POL III bound to SINE elements is transcriptionally active, we established a mESC line that enables conditional depletion of GTF3C1, the central subunit of the POL III recruitment factor TFIIIC ^37,38^. Treatment with dTAG13 strongly reduced GTF3C1 levels within 1 hour, with further reductions after prolonged treatment for 4 and 24 hours (h) (Figure 1D). Although some protein remained after 24 h, POL III ChIP-seq analysis revealed a near-complete loss of POL III binding at TFIIIC-dependent loci after 24 h of GTF3C1 depletion (Figures 1E). In contrast, POL III levels at TFIIIC-independent U6 and 7SK promoters remained unchanged, confirming the specificity of the depletion system (Figure 1E). Quantification of RNA-seq reads mapping to intergenic SINE insertions indicated that transcripts from the SINE B2 family are by far the most abundant in mESCs (Figure S1G). Notably, the evolutionary old families of B4, ID, and MIR elements are not bound by POL III. We nevertheless detected RNA reads mapping to these insertions. Likely, these originate from non-specific, spurious Pol II transcription events. This notion is supported by the observation that only transcripts of young SINE B2 elements, which are bound by POL III, were reduced upon TFIIIC depletion (Figures 1F, S1G). RNA levels of the other families, including evolutionary older SINE B2s that are not bound by POL III, did not react. Interestingly, RNA levels of young B1 elements were rather low and did not decrease upon TFIIIC depletion, despite some insertions being POL III bound (Figures 1A, S1G). As only 0.36 % of young SINE B1 insertions are POL III bound, we might be unable to specifically detect their transcriptional output in bulk assays considering the uniform background.

To strengthen the RNA-seq quantification of highly repetitive SINE sequences with an orthogonal approach, we performed Northern blotting with probes specific to B1 and B2 families of SINEs (Figure 1G, S1H). This confirmed that B2 RNAs are strongly reduced upon GTF3C1 depletion, suggesting that they are bona fide POL III transcripts. Consistent with RNA-seq, SINE B1 transcripts were not strongly decreasing after 24 h of GTF3C1 depletion. Of note, expression of Pol II-transcribed genes changed minimally, if at all, 4 h post GTF3C1 depletion. After 24 h, however, we observed extensive expression changes for hundreds of genes. We speculate this reflects a pleiotropic stress response caused by prolonged impairment of POL III transcription (Figure 1H).

In summary, using genetic and genomics approaches we show that evolutionary old SINE families, such as B4, ID, or MIR elements, are not bound by the POL III machinery. Few (0.36 %) copies of the younger SINE B1/Alu subfamilies Mm, Mus_1, and Mus_2 are bound by POL III but do not produce detectable amounts of GTF3C1-dependent RNA when quantified by RNA-seq or Northern blotting. However, copies belonging to elements of the young SINE B2 subfamilies Mm1a, Mm1t and Mm2 are bound by RNA POL III 3.5-fold more often (1.28 %) and produce detectable RNA. This establishes Mm1a, Mm1t, and Mm2 SINE B2 elements in mESCs as a suitable paradigm to study the regulation of POL III transcription of non-autonomous retrotransposons.

### SINE B2 elements are directly repressed by ChAHP

Initial studies had employed DNA or H3K9 methylation inhibitors to assess whether SINEs might be repressed by DNA methylation or heterochromatin formation ^31,32,39^. Although such treatment led to increased SINE transcript levels, a causal relationship between DNA or H3K9 methylation and SINE repression has not been established. To revisit this question, we analyzed published RNA-seq data sets from genetic depletion of either DNA methylation ^40^ or H3K9me3 ^41,42^ in mESCs. Consistent with previous reports, ERV and L1 transposons showed dependencies on H3K9me3 and DNA methylation. However, neither the knockout of all three DNA methyltransferases, nor *Su(var)39h1/2* double-knock out or SETDB1 depletion (H3K9 methylation) did affect SINE expression (Figure 2A). This raises the question of whether SINE expression is regulated by DNA or H3K9 methylation at all.

**Figure 2.**
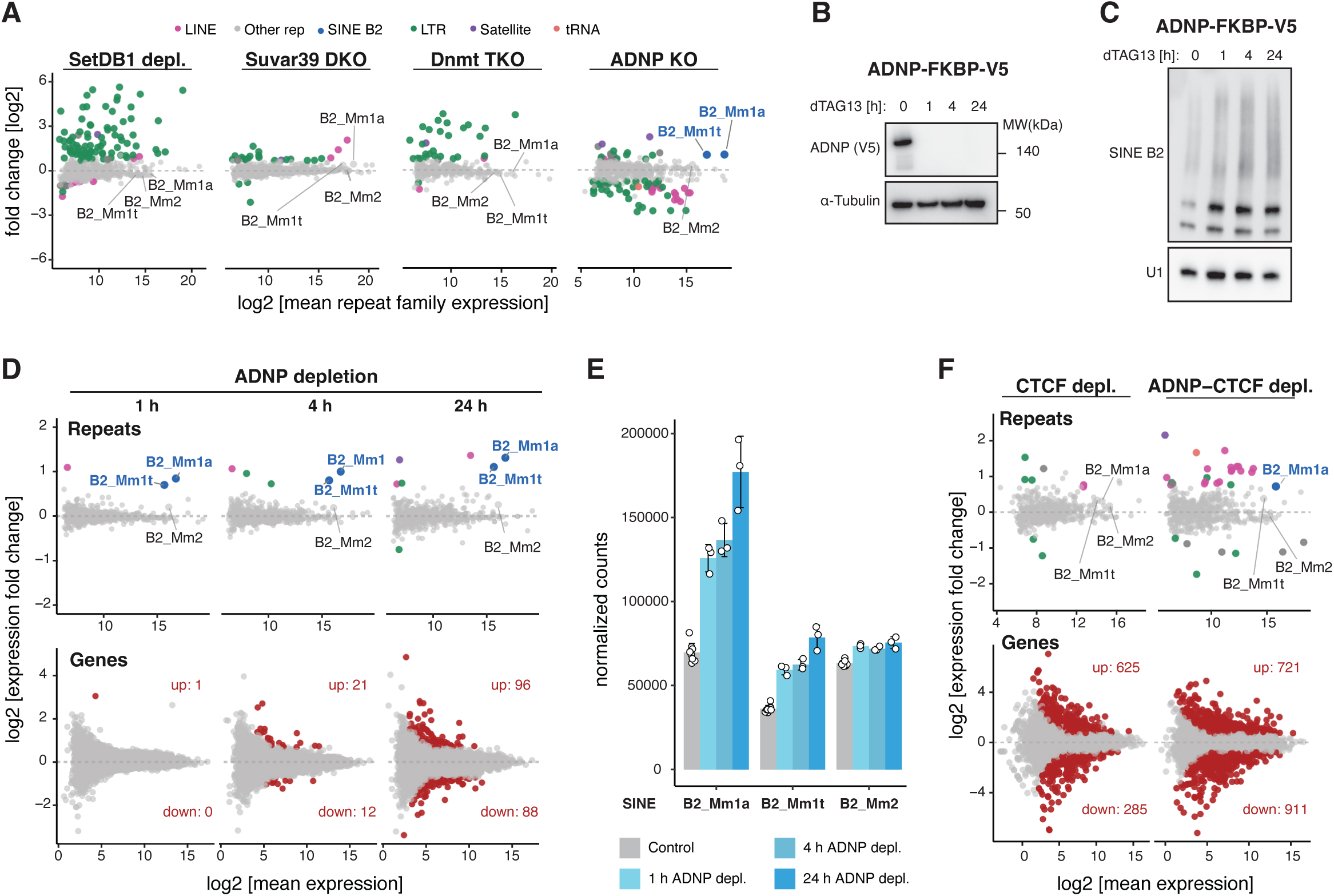
SINE B2 elements are directly repressed by ChAHP. **(A)** MA plots depicting differentially expressed repeat elements in ES cells with 48h SetDB1 depletion, Suvar39 DKO, Dnmt TKO, and ADNP KO. Significantly regulated repeats are highlighted in color, SINE B2 subfamilies are labeled. **(B)** Immunoblot showing levels ADNP-FKBP12-V5 fusion protein after treatment with 250nM of dTAG13 for the indicated times. α-Tubulin serves as loading control. **(C)** Northern blot analysis of SINE-B2 RNA expression following ADNP degradation. U1 snRNA serves as loading control. **(D)** MA plots showing differential gene expression after 1h, 4h, and 24h of ADNP depletion. Significantly regulated repeats and genes (padj < 0.05) are highlighted in color, SINE B2 subfamilies are labeled. **(E)** Quantification of normalized SINE-B2 subfamily transcript levels in control and ADNP depletion conditions at indicated time points. Data represent mean ± SD from three independent replicates. **(F)** MA plots comparing differential repeat and gene expression following 24h of CTCF depletion (left) versus combined ADNP-CTCF depletion (right). Significantly regulated repeats and genes (padj < 0.05) are highlighted in color, SINE B2 subfamilies are labeled.

In contrast, ablation of the ChAHP complex by knocking out *Adnp* resulted in increased SINE B2 RNA levels ^33,34,42,43^ (Figure 2A). To further test if ChAHP is directly repressing SINE B2 expression, we depleted ADNP conditionally. ADNP protein levels were strongly reduced within one hour of dTAG13 treatment and remained undetectable by Western blot after 24 h (Figure 2B). Northern blotting and RNA-seq of ADNP depleted cells showed that SINE B2 RNAs are upregulated as early as one hour after ADNP depletion, with the strongest effect observed for Mm1a and Mm1t subfamilies (Figure 2C-E). SINE B1 transcript levels remained unchanged, indicating that ChAHP-mediated repression is specific to active SINE B2 elements (Figure S2A-B). Importantly, the up-regulation of SINE B2 elements preceded any observable mis-regulation of RNA Pol II-transcribed genes, which start to be detected only after 4h of ADNP depletion and progressively increased at 24h (Figure 2D). This immediate effect on SINE B2 expression strongly supports a direct role of ChAHP in repressing these TEs.

We and others have previously reported that ChAHP counteracts CTCF binding at SINE B2 elements, which have acquired and spread CTCF binding motifs throughout the mouse genome ^7,33,43^. Given that CTCF is a multivalent chromatin regulator and its binding generates locally accessible chromatin with regularly spaced nucleosomes, both features of transcription-permissive chromatin, we depleted CTCF to test if its presence has an impact on SINE B2 expression. CTCF depletion alone had no effect on B2 RNA levels, whereas simultaneous depletion of ADNP and CTCF resulted in upregulation of Mm1a RNA, comparable to the effect observed upon ADNP depletion alone (Figure 2F, S2C-D). Therefore, we conclude that chromatin occupancy of CTCF at SINE B2s does not affect their expression. Collectively, these results support a direct role of ChAHP in repressing POL III transcription of evolutionary young SINE B2 retrotransposons, which is independent of ChAHP’s role in counteracting CTCF binding to these elements ^33,43^.

### ChAHP and TFIIIC bind SINE elements independently of each other

The most parsimonious mechanism by which ChAHP could repress transcription of SINEs is through steric hindrance of TFIIIC binding, the most upstream POL III general transcription factor (GTF) involved in initiating POL III transcription from type 2 promoters. Notably, studies in human cells have shown that ADNP physically interacts with TFIIIC and it was proposed that ADNP recruits TFIIIC to specific sites in the human genome ^44^.

To investigate whether these interactions occur between TFIIIC and ChAHP as intact complexes, we endogenously tagged the GTF3C1 subunit of TFIIIC and performed immunoprecipitation followed by mass spectrometry (IP-MS). Consistent with the observation in human cells, all subunits of the ChAHP complex co-purified with the 6 subunit TFIIIC complex (GTF3C1– GTF3C6). Reciprocally, ADNP purifications recovered all TFIIIC complex members (Figure 3A). To corroborate a direct physical interaction between ChAHP and TFIIIC, we purified both complexes and performed in-vitro pull-down experiments (Figure 3B). This revealed a direct interaction of intact ChAHP with TFIIIC, showing that ADNP can interact with TFIIIC as part of the ChAHP complex.

**Figure 3.**
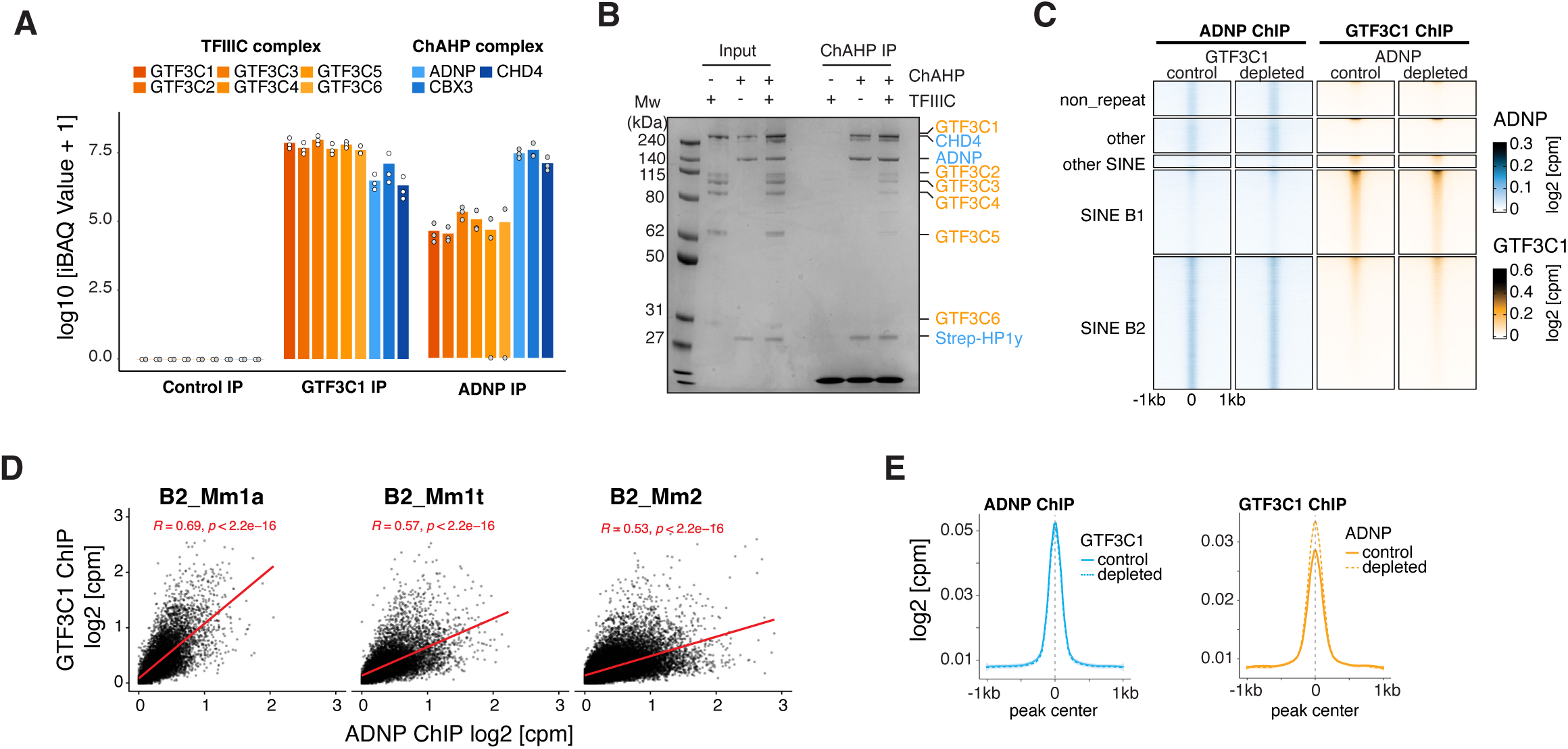
ChAHP and TFIIIC bind SINE B1 and B2 elements independently of each other. **(A)** Quantification of mass spectrometry analysis (iBAQ values) after immunoprecipitation of GTF3C1 (open circles) or ADNP (grey circles) compared to untagged controls. Subunits of TFIIIC (orange) and ChAHP (blue) are indicated. **(B)** Coomassie stained SDS-PAGE gel showing *in vitro* pulldown of ChAHP with TFIIIC. Complexes alone serve as control for background binding. Molecular weight markers (left) and protein identities (right) are indicated. **(C)** Heatmaps showing ADNP and GTF3C1 ChIP-seq signal in control and depletion conditions across different genomic features. **(D)** Scatter plots showing correlation between ADNP and GTF3C1 ChIP-seq enrichment at individual loci of SINE-B2 subfamilies. Pearson correlation coefficients (R) and p-values are indicated. **(E)** Meta-profiles of ADNP and GTF3C1 ChIP-seq at SINE B2 peaks in control (blue and orange, respectively) and depletion conditions (dotted blue and yellow, respectively). Signal is plotted as log2 [cpm] relative to peak center with 1 kb flanking regions.

We next assessed whether ChAHP and TFIIIC affect each other’s occupancy on SINE elements by performing ChIP-seq with endogenously tagged GTF3C1 (TFIIIC) and ADNP (ChAHP). As expected, ADNP predominantly localized to SINE elements, with most peaks overlapping the B2 family ^33^ (Figure 3C). In addition to B2s, ADNP also showed low enrichment at B1 elements. GTF3C1 was enriched at POL III genes such as tRNAs as reported before ^26^. Among SINE families, we observed the strongest enrichment at B1 elements, followed by B2s (Figures 3C, S3B). Consistent with the observed physical interaction between ChAHP and TFIIIC, we saw a correlation of ADNP and GTF3C1 predominantly at evolutionary young B2 Mm1a and Mm1t subfamilies (Figure 3D).

To test a direct dependency of these two factors for chromatin binding, we depleted either GTF3C1 or ADNP and assayed ADNP or GTF3C1, respectively, for DNA binding (Figure S3A). Unlike reported in human cells ^44^ and despite their strong colocalization, depletion of ADNP or GTF3C1 revealed that the ChAHP and TFIIIC complexes can associate with their targets independent of each other (Figure 3C). Notably, when comparing chromatin occupancy at lowly divergent SINE B2 insertions, we observed a small increase of GTF3C1 binding, while ADNP binding remained unaffected upon GTF3C1 depletion (Figure 3E). This marginal increase might result from elevated POL III transcription in the absence of ChAHP (Figure 2D).

These results demonstrate that ADNP and TFIIIC maintain independent chromatin associations in mESCs and strongly argue against a model that ChAHP could repress POL III transcription by sterically hindering TFIIIC binding.

### ChAHP interferes with TFIIIB recruitment to repress POL III transcription

To comprehensively test at what step of POL III transcription initiation ChAHP interferes, we endogenously tagged subunits of all POL III GTFs in cells that allow conditional depletion of ADNP (Figure S4A). ChIP-seq of the POL III GTFs revealed robust enrichments at their respective targets (Figure S4B). As expected, SINE B1 and B2 insertions bound by ChAHP and/or GTF3C1 were only bound by GTFs of type 2 promoters (GTF3C1, BRF1) (Figures 4A-B).

**Figure 4.**
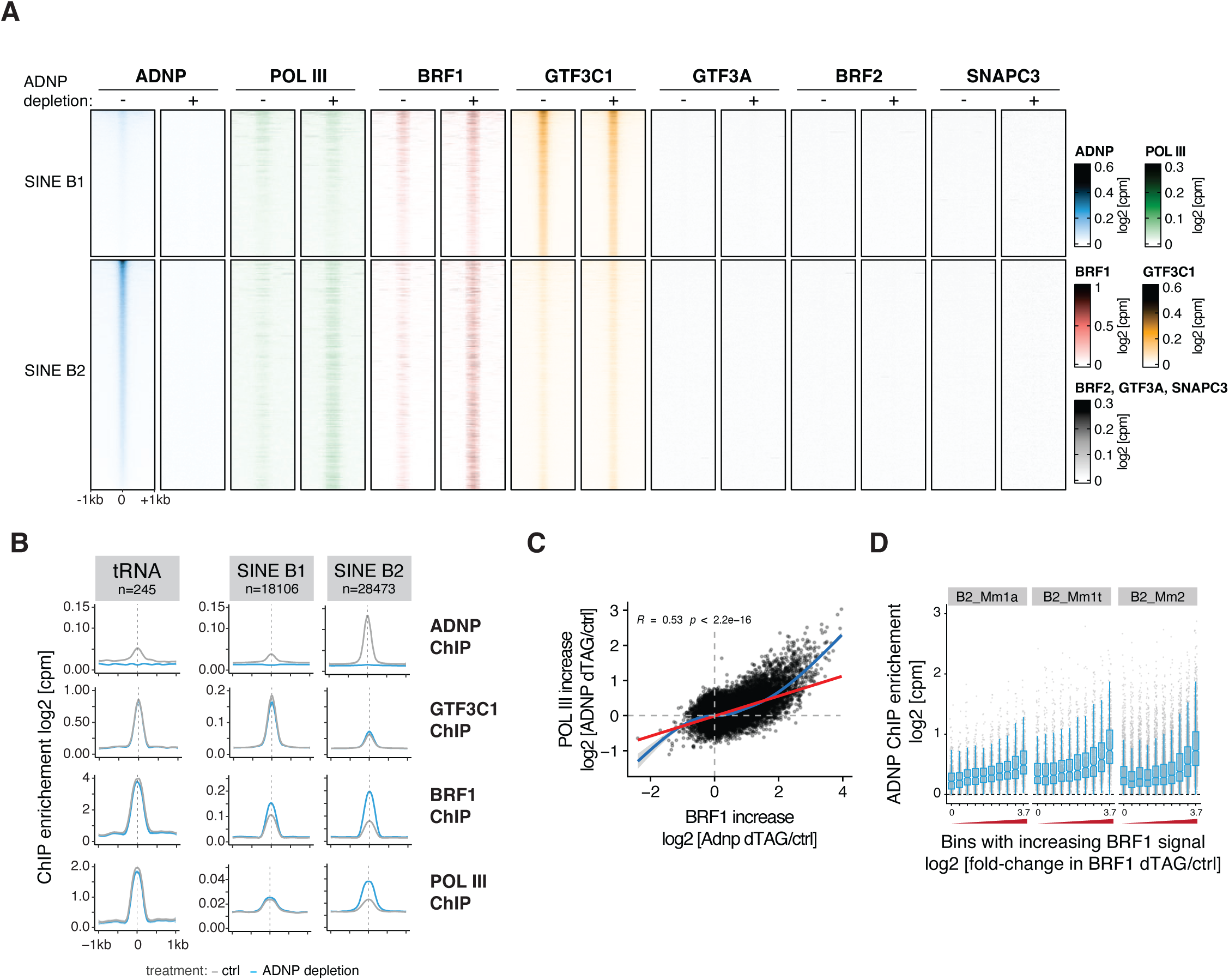
ChAHP interferes with TFIIIB recruitment to repress POL III transcription. **(A)** Heatmaps showing ChIP-seq enrichment for indicated factors at SINE B1 (upper panels) and SINE B2 (lower panels) elements in control (-) and ADNP-depleted (+) conditions. Color scales indicate log2 [cpm] values for each factor. **(B)** Meta-profiles of ChIP-seq signals for ADNP, GTF3C1, BRF1, and POL III across tRNAs, SINE B1s, and SINE B2s in control (grey) and 24 h ADNP depletion (blue) conditions. Profiles are centered on genomic elements with 1 kb flanking regions. **(C)** Correlation between BRF1 and POL III recruitment increases (log2 fold-change) upon ADNP depletion. Red line: linear regression with Pearson correlation coefficient (R) and p-value indicated. Blue line: locally weighted regression (loess). **(D)** Box plots showing ADNP ChIP enrichment (log2 [cpm]) at SINE B2 subfamilies stratified by BRF1 occupancy increase upon ADNP depletion. The x-axis represents bins of log2 fold-change in BRF1 occupancy from low to high.

Upon ADNP depletion, B1 elements, although weakly bound by ChAHP, acquired only marginally more BRF1 and POL III. In contrast, B2 elements, which are strongly enriched for ADNP in untreated cells, acquire high levels of BRF1 and POL III upon ADNP depletion, in agreement with their transcriptional upregulation. Importantly, other POL III-associated factors such as GTF3A, SNAPC, or BRF2 that are not implicated in type 2 promoter regulation remained absent at transcriptionally upregulated SINE B2s. Moreover, the levels of POL III GTFs at tRNA genes, which are also weakly bound by ADNP, remined unchanged (Figure 4B). To test if the same SINE B2 insertions that gain TFIIIB binding also show increased POL III binding and therefore are transcriptionally upregulated, we correlated the changes in ChIP enrichment upon ADNP depletion. Reassuringly, the elevated BRF1 ChIP signal strongly correlated with the increase in POL III, supporting the conclusion that the increase in TFIIIB occupancy upon ChAHP depletion is causative for the enhanced POL III transcription at SINE B2s (Figure 4C).

Finally, we investigated whether ChAHP occupancy could predict the degree of repression, or conversely, the extent of increased BRF1 and POL III enrichment in the absence of ChAHP. When binning SINE B2 transposon insertions according to their increase in BRF1 enrichment upon ADNP depletion, we observed that insertions with higher ADNP levels exhibited stronger increases in BRF1 ChIP signal upon ADNP depletion (Figure 4D). In agreement with the SINE B2 expression data, this trend was particularly evident at Mm1a and Mm1t elements. Since these are the evolutionary youngest elements and ChAHP preferentially binds to lowly divergent copies ^33^, we assessed whether the transcriptional upregulation also correlates with sequence divergence, i.e. evolutionary age. This is indeed the case; additionally, the upregulated copies exhibit the highest scores for A-box and B-box motifs (Figure S4G-H).

Collectively, our results demonstrate that ChAHP specifically interferes with TFIIIB recruitment at evolutionarily young SINE B2 elements, revealing a targeted mechanism for silencing POL III-transcribed transposable elements capable of retro-transposition in the mouse genome.

## DISCUSSION

Our findings establish the ChAHP complex as a direct repressor of POL III transcription specifically at SINE transposable elements. This conclusion is based on the following key findings from our comprehensive analysis of POL III GTFs and ChAHP: (1) ChAHP is specifically targeting young SINE B1 and B2 insertions, (2) B2 transcripts are upregulated within one hour of ADNP depletion, whereas changes in protein coding genes are only detectable after prolonged depletion, (3) TFIIIC, the most upstream GTF for POL III transcription, is already pre-bound to many inactive and upregulated loci and does only marginally increase while (4) TFIIIB and POL III levels are strongly elevated upon ADNP depletion, and (5) other POL III genes, such as tRNAs with similar promoter types as SINE B2s, are not affected by loss of ADNP.

The fact that ChAHP acts on TFIIIB is particularly intriguing, as TFIIIB is not only a recruitment factor but also plays essential roles in pre-initiation complex formation ^19^. Moreover, POL III’s ability to achieve high transcriptional output is thought to depend on facilitated reinitiation, a specialized transcription mode that bypasses the need for TFIIIC but likely requires TFIIIB ^45–47^. By targeting TFIIIB recruitment, ChAHP effectively blocks both the initiation of transcription and this putative reinitiation pathway, providing a potent mechanism for silencing SINE elements.

On a mechanistic level, we propose two possible models for how ChAHP might counteract TFIIIB recruitment: Either, the binding of the multi-subunit ChAHP complex sterically hinders TFIIIB recruitment. Alternatively, the ATPase activity of CHD4 is employed to reposition nucleosomes and rendering the TFIIIB binding motif inaccessible, or to continuously expel TFIIIB from SINE B2 promoters. In a related preprint, we show that the catalytic activity of CHD4 within the ChAHP complex is essential for SINE B2 repression, strongly supporting the latter model (Ahel *et al.* 2025. Remodeling Activity of ChAHP Restricts Transcription Factor Access to Chromatin. *bioRxiv*). Interestingly, earlier studies have revealed that transcription initiation by POL III is sensitivity to nucleosome positioning over promoter elements, adding further support for such a mode of action^48,49^.

Importantly, the mode of silencing revealed in our study operates distinctly from canonical transposon silencing mechanisms, which typically rely on heterochromatin formation or DNA methylation ^50^. Such chromatin-based mechanisms often involve spreading of repressive marks, leading to the establishment of heterochromatic domains spanning several kilobases. Whereas this is well-suited to repress long TEs, such as LINEs and ERVs that span 4-10kb, it may be less effective for SINE elements, which are only 80-300bp in length and encompass just 1-2 nucleosomes. Moreover, due to their exceptionally high copy number, chromatin-based SINE silencing could lead to unwanted off-target repression of nearby transcription units and other regions across the genome. In contrast, ChAHP’s sequence specific nucleosome remodeling activity provides a locally restricted mechanism for silencing POL III transcription. Intriguingly, Fpt1 was recently identified as a *Saccharomyces cerevisiae*-specific POL III repressor that can mediate tRNA silencing under certain metabolic conditions via chromatin-based repression of POL III transcription ^51^. Although the underlying molecular mechanism is unknown, increased Fpt1 binding at tRNA genes under repressive conditions coincides with enrichment of chromatin remodelers such as INO80. This raises the possibility that Fpt1 and ChAHP utilize similar molecular strategies for inhibiting POL III transcription.

Of note, the sequence-specific repression of POL III by ChAHP fundamentally differs from the mechanism employed by Maf1, the sole other known POL III repressor in mammals. MAF1 is phosphorylated and thereby restricted to the cytoplasm under normal growth conditions. Upon nutrient deprivation, mTOR signaling decreases MAF1 phosphorylation. This allows MAF1 to enter the nucleus and bind POL III at a position that prevents interaction with TFIIIB, thereby preventing proper transcriptional initiation at all POL III genes ^52^. Yet, despite employing distinct mechanisms, both ChAHP and MAF1 converge on interfering with TFIIIB, which is essential for assembling a transcription-competent POL III complex.

### Limitations of the study

It is important to note that assigning transposon-derived transcripts to specific loci is inherently difficult due to their repetitive nature. This challenge is particularly pronounced for SINEs, given their extremely high copy number and frequent localization within introns of protein-coding genes. Although we employed multiple orthogonal approaches to identify genuine POL III-driven transcription, we cannot fully exclude the possibility that some RNA-seq signals reflect Pol II-derived transcripts.

This study was conducted in mESCs, which are amenable to genome engineering and well-suited for dissecting chromatin regulatory processes. We demonstrate that SINE elements exhibit basal levels of transcription that is attenuated by the ChAHP complex, making mESCs an ideal system for studying SINE regulation. Whether the regulatory principles worked out in this study extend to other cell types or developmental stages remains to be determined. Further, our study does not address the functional significance of SINE transcripts or the physiological consequences of their upregulation following ADNP depletion. Given proposed roles for SINE RNAs in modulating RNA polymerase II activity ^53^, it is possible that ChAHP indirectly influences Pol II-driven transcription of protein-coding genes, beyond its established role in regulating 3D chromatin organization ^33,43^.

## Supporting information

Sequences of gRNA oligos and homolgy donor constructs used for Cas9-mediated genome engineering.

Cell lines used in this study.

## RESOURCE AVAILABILITY

### Lead contact

Further information and requests for resources and reagents should be directed to and will be fulfilled by the lead contact, Marc Bühler.

### Materials availability

mESC lines and plasmids are available upon request.

#### Data and code availability

Sequencing data from ChIP-seq and mRNA-seq have been deposited at GEO (accession number GSE297837). The mass spectrometry proteomics data have been deposited to the ProteomeXchange Consortium via the PRIDE^54^ partner repository with the dataset identifier PXD064515. Accession numbers of published data sets used in this study are listed in the key resources table.

Original code has been made available on GitHub (https://github.com/xxxmichixxx/ADNP_POL3).

## ACKNOWLEDGEMENTS

We thank all members of the Bühler lab for discussions, comments on the manuscript and constant support. We thank all members of the FMI genomics, proteomics and structural biology platforms for their excellent support, especially Daniel Hess for Mass spectrometry analysis. We thank Hubertus Kohler for cell sorting. We are thankful to Si Hoon Park for advice on insect cell culture and virus production. We are grateful to Lisa Baumgartner, Merle Skribbe, Dominik Handler, Maya Voichek, Changwei Yu and Lucas Kaaij for their comments on the manuscript. This work was supported by the Novartis Research Foundation and the Swiss National Science Foundation (SNSF; grant 10.001.858). Jakob Schnabl-Baumgartner was supported by an EMBO Postdoctoral Fellowship (ALTF 705-2021).

## Author contributions

Conceptualization: J.S-B., F.M., M.B; Data curation: F.M., M.S; Formal analysis: J.S-B., F.M., M.S; Funding acquisition: J.S-B., M.B; Investigation: J.S-B., F.M., J.A, J.S; Methodology: J.S-B., F.M.; Project administration: J.S-B., F.M., M.B; Resources: J.S-B., F.M., M.S, J.S, Y.S; Software: M.S; Supervision: M.B; Validation: J.S-B., F.M.; Visualization: J.S-B., F.M., M.S; Writing – original draft: J.S-B., F.M.; Writing – review & editing: J.S-B., F.M., M.S, J.A, J.S, M.B

## Declaration of generative AI and AI-assisted technologies in the writing process

During the preparation of this work the authors used Claude Sonnet 3.7 to improve the readability and language of the manuscript. After using this tool, the authors reviewed and edited the content as needed and take full responsibility for the content of the published article.

---

**Figure S1.**
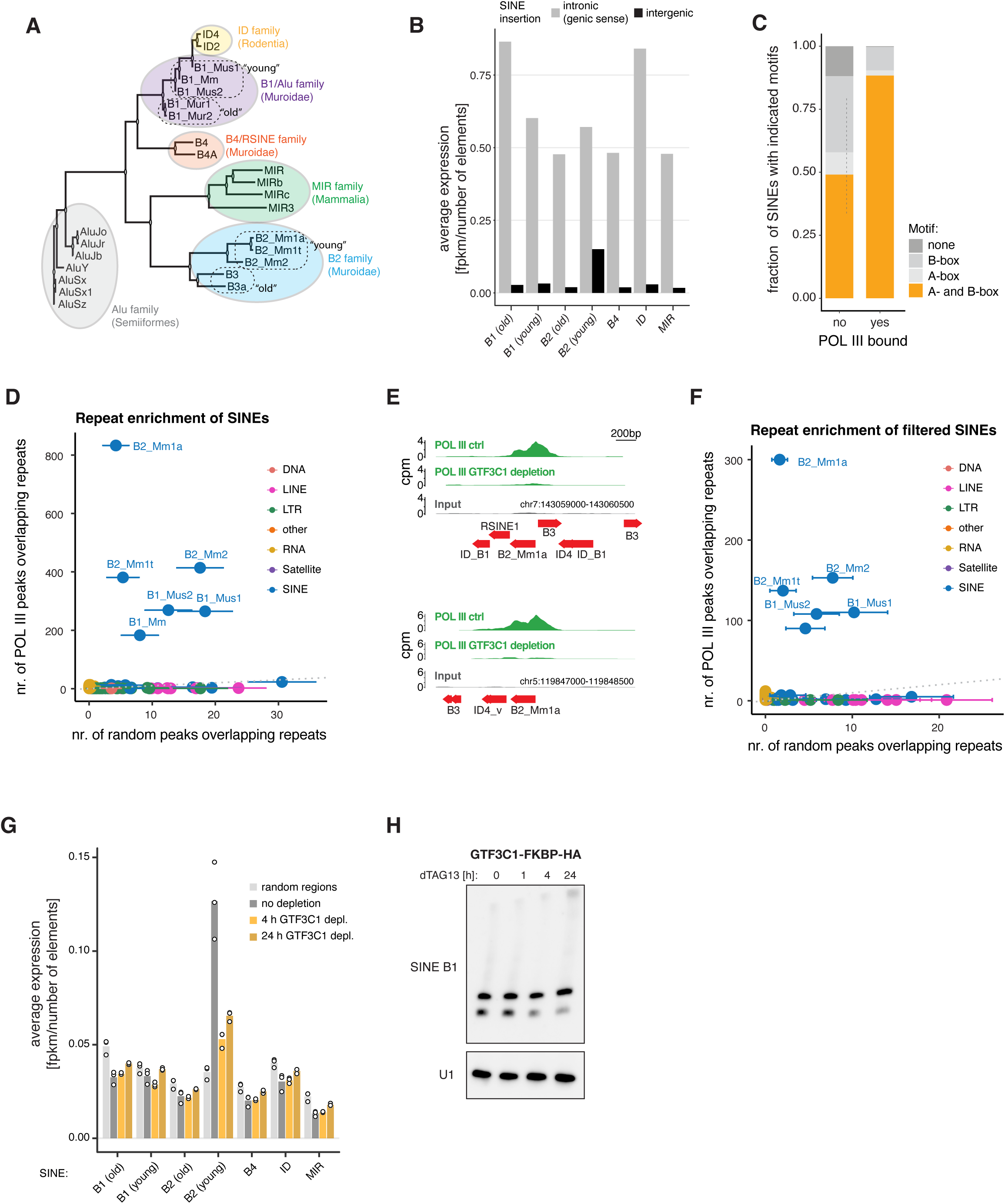
SINE B2 elements show the strongest signatures of POL III transcription, related to Figure 1. **(A)** Phylogenetic tree of the five most abundant SINE families and their subfamilies in mouse and human. Age grouping for B1s and B2s indicated by dashed circles. **(B)** Average expression (fpkm/number of elements) of different SINE subfamilies in genic versus intergenic regions. For B1/B2 young/old, see grouping in S1A. **(C)** Distribution of motifs (none, B-box only, A-box only, both A- and B-box) in SINEs with and without POL III enrichment. **(D)** Enrichment analysis of repeat subfamilies in POL III ChIP-seq peaks compared to random genomic regions. The number of repeats of a given subfamily that overlap a POL III peak is shown on the y-axis. The x-axis shows the mean and standard deviation of the number of repeats of a given subfamily that overlap 50 sets of random peaks of the same width and number as observed POL III peaks. **(E)** Genome browser tracks showing POL III ChIP-seq signal at two representative genomic loci containing SINE B2 elements. **(F)** Enrichment analysis of repeat subfamilies in POL III ChIP-seq peaks compared to random genomic regions as in **(C)**, but using only those repeats that do not neighbor another POL III transcription unit within 200b. **(G)** Average expression levels (fpkm/number of elements) of different SINE subfamilies in control and following 4 h or 24 h of GTF3C1 depletion. For B1/B2 young/old, see grouping in S1A. **(H)** Northern blot analysis showing expression of SINE B1 RNA following GTF3C1 depletion for the indicated times. U1 snRNA serves as loading control.

**Figure S2.**
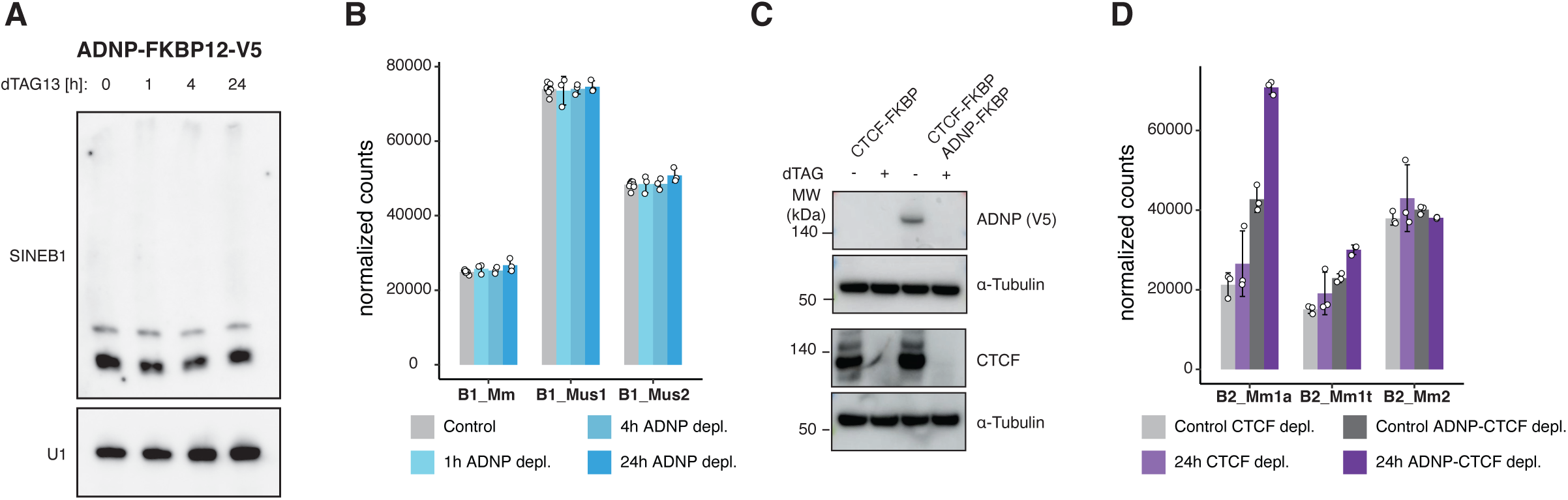
SINE B1 expression is not affected after ADNP depletion, related to Figure 2. **(A)** Northern blot analysis of SINE B1 RNA expression during time course of ADNP depletion. U1 snRNA shown as loading control. **(B)** Quantification of normalized SINE B1 subfamily transcript levels (B1_Mm, B1_Mus1, B1_Mus2) in control cells and following ADNP depletion. Data represent mean ± SD from three independent experiments. **(C)** Immunoblot analysis validating protein depletion in the indicated cell lines following dTAG treatment. Blots were probed with antibodies against ADNP (V5-tag), CTCF, and α-Tubulin (loading control). **(D)** Normalized expression of SINE B2 subfamily transcripts comparing CTCF depletion alone versus ADNP-CTCF co-depletion. Data represent mean ± SD from three independent experiments.

**Figure S3.**
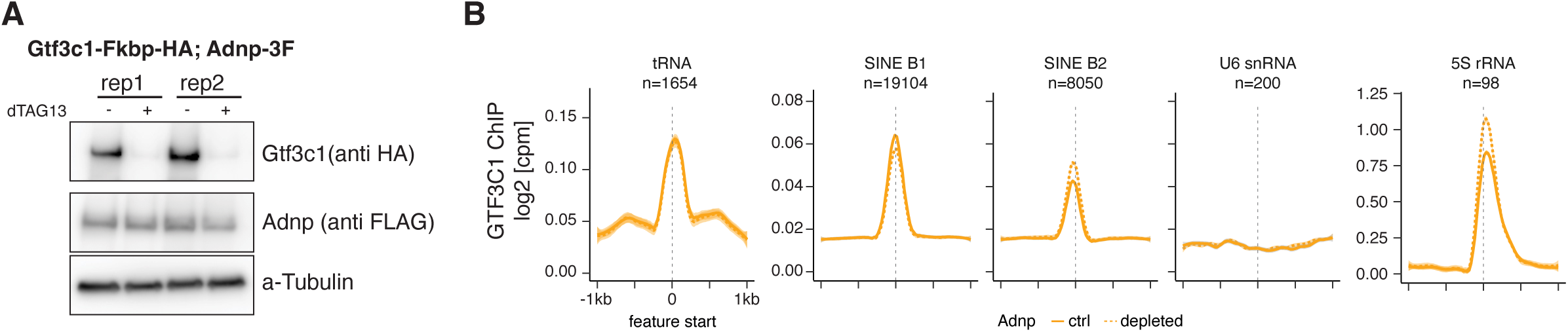
TFIIIC binding at POL III transcribed loci is not affected upon ADNP depletion, related to Figure 3. **(A)** Immunoblot analysis of GTF3C1-Fkbp-HA ADNP-3xFLAG cell lines (two independent replicates) treated with or without 250nM dTAG13. Blots were probed for GTF3C1 (anti HA), ADNP (anti-FLAG), and α-Tubulin (loading control). **(B)** Meta-profiles of GTF3C1 ChIP-seq at POL III transcribed loci in control (grey) and 24 h ADNP depleted conditions (orange). Signal is plotted as log2 [cpm] relative to feature start with 1 kb flanking regions.

**Figure S4.**
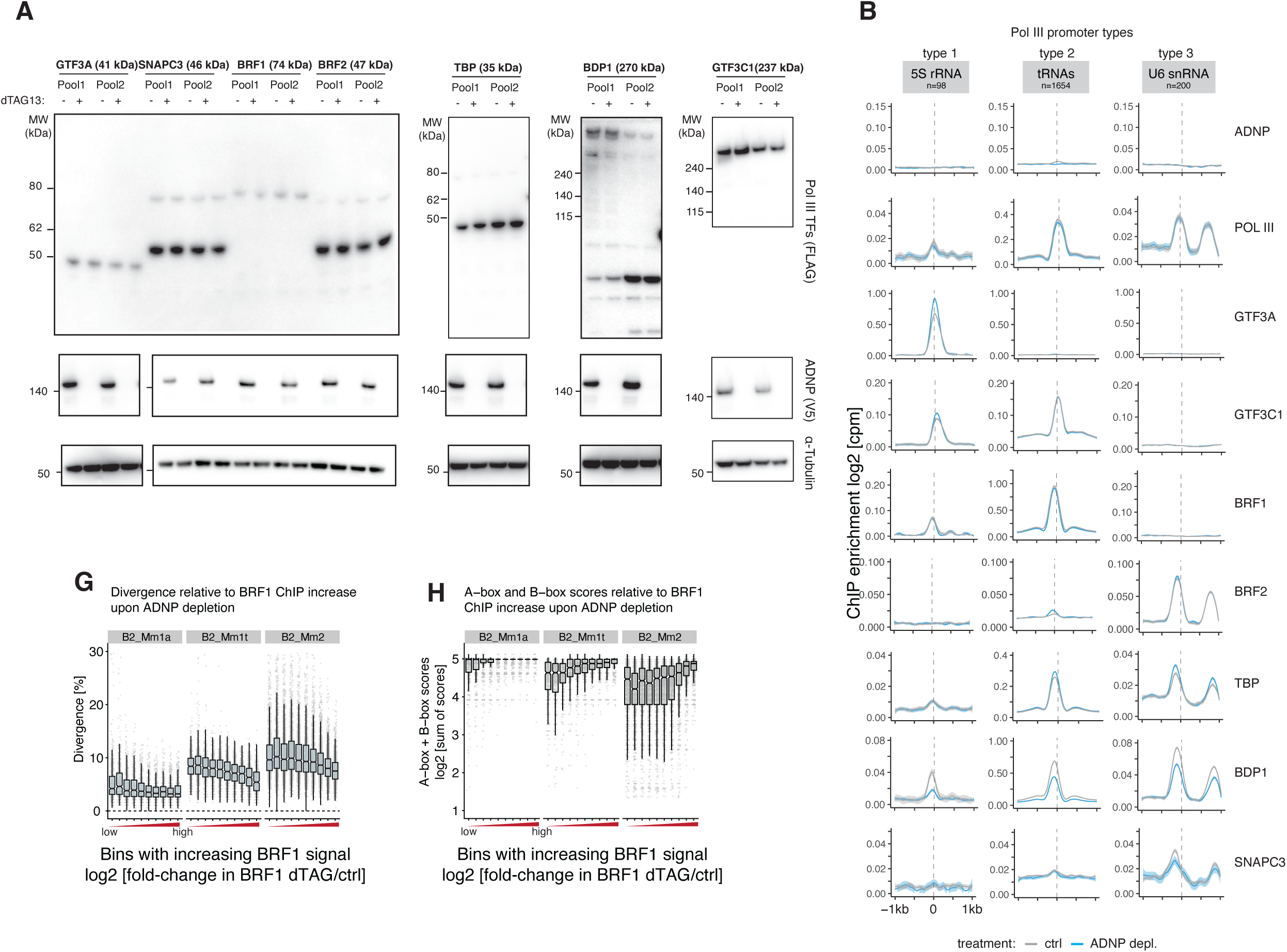
Endogenously tagged POL III GTFs bind to their respective targets, related to Figure 4. **(A)** Immunoblot analysis validating expression levels of POL III transcription factors in ADNP-FKBP cells in the absence (-) or presence (+) of 250nM dTAG13. Two independent cell pools were analyzed for each factor. Molecular weights are indicated. α-Tubulin serves as loading control. **(B)** Meta-profiles of ChIP-seq signals (log2 [cpm]) for RNA POL III, BRF2, SNAPC3, and GTF3A across type 1 (5S rRNAs, n=98), type 2 (tRNAs, n=1,654), and type 3 (U6 snRNAs, n=200) POL III genes in control (grey) and ADNP-depleted (blue) conditions. **(G)** Box plots showing sequence divergence (%) of SINE B2 subfamilies stratified by BRF1 occupancy increase following ADNP depletion. **(H)** Box plots showing combined A-box and B-box motif scores [log2] at SINE B2 subfamilies stratified by BRF1 occupancy increase following ADNP depletion.

## METHODS

### Mouse embryonic stem cell culture

Mouse embryonic stem cells (129 × C57BL/6 background) with BirA and Cre insertions in the Rosa26 locus were used as wildtype background ^34,55^. All cell lines were derived from the same parental clone to ensure isogenic cell lines. Cells were maintained at 37°C, 5% CO2 in mESC medium containing DMEM (GIBCO), 15% fetal bovine serum (FBS; GIBCO), 1 × non-essential amino acids (GIBCO), 1 mM sodium pyruvate (GIBCO), 2 mM l-glutamine (GIBCO), 0.1 mM 2-mercaptoethanol (Sigma), 50 mg/ml penicillin, 80 mg/ml streptomycin, 3 μM GSK3 inhibitor (STEMCELL Technologies), 10 μM MEK inhibitor (Tocris, PD0325901), and homemade LIF.

### SF9 insect cell culture

SF9 cells were obtained from ThermoFisher cultured in SF-4 Baculo Express ICM media (BioConcept) at 27°C in a shaking incubator at 120 RPM. Cells were split upon reaching a density of around 4 million cells per mL by dilution with fresh media to 0.5 million cells per mL.

### Mouse embryonic stem cell engineering

For modification of the endogenous locus of ADNP in mESCs, TALENs were used, while CRISPR-Cas9 mediated endogenous tagging strategy was used for other targets. The specific N- or C-terminal tagging approach and introduced tags are indicated in Table S1.

For both approaches, homology-directed repair (HDR) templates were assembled in pBluescript vectors with 200-400 bp long homology arms surrounding the cut-sites. When applicable, synonymous mutations at the PAM sequences of the guide RNAs were introduced to prevent cutting of the repaired allele. Guide RNA-containing Cas9 expression vectors were cloned using two annealed single-stranded DNA oligonucleotides (IDT) with the sequences provided in Table S1.

500,000 cells were seeded into a 6-well plate and transfected using Lipofectamine 3000 reagent (Invitrogen) with 187 μL Opti-MEM (Thermo Fisher), 3 μL P3000 reagent, and 4.5 μL Lipofectamine 3000 for 1.5 μg of plasmid DNA according to the manufacturer’s instructions. When applicable, 24 h after transfection, selection with 0.5 μg/mL puromycin (Invitrogen) was performed for 24-32 hours.

For generating clonal cell lines, single cells were sorted into 96-well plates and further characterized after approximately 1 week using PCR, Sanger sequencing, and Western blotting. For POL III GTF pools, the HDR template contained a P2A sequence followed by mNeonGreen, which allowed FACS sorting of fluorescent cells. The sorting was repeated 1-2 times to achieve high purity and enrichment for homozygously tagged cells. Lines were validated by Western Blotting and partially IP-MS experiments

### ChIP sequencing

#### Chromatin immunoprecipitation (ChIP)

ChIP sequencing was performed with at least two replicates per condition. 30 μL of a 1:1 mixture of Protein A/G Dynabeads (Invitrogen) was pre-coupled with 3 μL of FLAG antibody (Sigma-Aldrich, FLAG M2 mouse mAb), V5 antibody (Invitrogen, SV5-Pk1 mouse mAb), or POL III antibody (Cell Signaling Technology, POLR3A (D5Y2D) Rabbit mAb) for 1 hour at room temperature. Supernatant was discarded and antibody coupled beads were resuspended in 30 µl 1× ChIP buffer (0.01% SDS, 1% Triton X-100, 1.2 mM EDTA, 16.7 mM Tris-HCl pH 8.0, 167 mM NaCl) and stored on ice until use.

A total of 10 million cells were crosslinked for 8 minutes at room temperature in 10 mL PBS supplemented with 1% formaldehyde (Sigma). The reaction was quenched by adding glycine to a final concentration of 0.125 M. Cells were pelleted by centrifugation at 500 × g for 3 minutes at room temperature and supernatant was completely removed. The cells were lysed in 10 mL of buffer A (50 mM HEPES pH 8.0, 140 mM NaCl, 1 mM EDTA, 10% glycerol, 0.5% NP-40, 0.25% Triton X-100) on ice for 10 minutes. After centrifugation, the pellet was resuspended in 5 mL of buffer B (10 mM Tris pH 8.0, 1 mM EDTA, 0.5 mM EGTA, 200 mM NaCl) and incubated on ice for 5 minutes. Nuclei were pelleted by centrifugation at 500 × g for 3 minutes at 4°C, resuspended in 150 μL of buffer C (50 mM Tris pH 8.0, 5 mM EDTA, 1% SDS, 100 mM NaCl), and incubated on ice for 10 minutes. Chromatin was sheared for 30 minutes in a PIXUL 96-well sonicator (Active Motif) using 2 wells per sample (i.e. 5 mio cells/well) and the following settings: 50 N Pulse, 1 ms Pulse Duration 1 kHz PRF, 20 Hz Burst Rate. After shearing, the corresponding wells for each sample were pooled and 1x ChIP buffer was added to a final volume of 2ml. The chromatin was transferred to fresh tubes and centrifuged at 13,000 × g for 10 minutes at 4°C. Per sample, 30 μL of 1:1 pre-coupled beads were added to the chromatin, and the mixture was incubated with rotation for 4 hours at 4°C. The ChIP samples were washed four times with low salt buffer (10 mM Tris-HCl pH 8.0, 1 mM EDTA pH 8.0, 140 mM NaCl, 1% Triton X-100, 0.1% SDS, 0.1% sodium deoxycholate), twice with high salt buffer (same as low salt but with 500 mM NaCl), twice with LiCl buffer (10 mM Tris-HCl pH 8.0, 1 mM EDTA pH 8.0, 250 mM LiCl, 0.5% NP-40, 0.5% sodium deoxycholate), and once with TE+ buffer (10 mM Tris-HCl pH 8.0, 1 mM EDTA, 50 mM NaCl). All buffers were cooled on ice before washing. During the final wash, beads were transferred to fresh tubes, and the wash buffer was completely removed. DNA was eluted twice by incubation at 65°C with shaking using 75 μL of elution buffer (10 mM Tris-HCl pH 8.0, 1 mM EDTA pH 8.0, 150 mM NaCl, 1% SDS), eluates were pooled. Input samples were adjusted to a total volume of 150 μL with elution buffer and processed in parallel to ChIP samples. 2 μL of RNase A (20 μg/μL) was added to each ChIP or Input sample, followed by incubation for 1 hour at 37°C. Proteinase K (2 μL, 20 mg/mL) was added, and samples were incubated for 2 hours at 55°C and reverse cross-linked for 6 hours at 65°C. Then, 9 μL of 5 M NaCl, 30 μL of AMPure XP beads (Beckman Coulter), and 190 μL of isopropanol were added. Samples were vigorously mixed, incubated at room temperature for 10 minutes, and the beads were collected on a magnetic rack. The beads were washed twice with 80% ethanol, and DNA was eluted in 30 μL of 10 mM Tris-HCl pH 8.0 for 5 minutes at 37°C.

#### Library preparation and sequencing

For library preparation, 25 μL of ChIP DNA or 10 ng of input DNA was used with the NEBNext Ultra II Library Prep Kit for Illumina (New England Biolabs), following the manufacturer’s protocol but with reactions scaled down to half-volume. Libraries were sequenced on the NovaSeq 6000 platform (Illumina) with paired-end reads (2 × 56 bp).

### Cell lysis and Western blotting

After trypsinization, 500,000 cells were harvested by centrifugation and washed once with 1× PBS. The cell pellet was resuspended in 40 μL of TMNS buffer (10 mM Tris pH 8, 1 mM MgCl₂, 100 mM NaCl, 1% SDS) supplemented with 0.2 μL Turbonuclease (Merck) and 0.4 μL 100× proteinase inhibitor cocktail (Thermo Fisher Scientific) and incubated at room temperature for 2 min. After vortexing the lysate for 5 s, the lysate was kept on ice until further use. 10 μL of 5× SDS loading dye were added followed by denaturation at 75°C for 5 min. For protein separation, 10 μL of lysate (equivalent to 100,000 cells) was loaded onto a Tris-Acetate gel (Thermo Fisher Scientific) and electrophoresed at 200 V for 30 min. Proteins were transferred to a Immobilon-P PVDF membrane (Millipore) using the Bio-Rad Trans-Blot Turbo transfer system. After blocking with 4% milk in TBST (1× TBS, 0.05% Tween-20) for 30 min at room temperature, the membrane was incubated with primary antibodies (FLAG antibody (Sigma-Aldrich, FLAG M2 mouse mAb, 1:2000), V5 antibody (Invitrogen, SV5-Pk1 mouse mAb, 1:2000), Anti-HA (Sigma-Aldrich, mouse mAb, 1:2000), anti-alpha-tublin (Gull lab, mouse mAb, 1:8000)), overnight at 4°C. The membrane was then washed 3 times for 15 minutes with TBST and incubated with HRP-conjugated secondary antibodies (goat anti-mouse, Jackson immunology) diluted 1:4,000 in 4% milk in TBST for 1 h at room temperature. After washing 4 times for 5 min with TBST, the chemiluminescence signal was developed using the Immobilon Chemiluminescent HRP Substrate (Millipore) and recorded on the Amersham Imager 680 (Cytiva).

### RNA extraction and Northern blotting

200,000 cells were seeded 24 h before RNA extraction and treated with dTAG13 or DMSO for indicated times. RNA was extracted using TRIzol reagent (Ambion) according to manufacturer’s instructions. 200 ng of RNA was diluted in 5 μL of water and mixed with 2× Novex TBE-Urea sample buffer (Thermo Fisher) and denatured at 95°C for 3 min before being placed immediately on ice. RNA was separated on 6% Novex Bolt Mini gel (Invitrogen) with 1× TBE buffer pre-heated to 50°C for 90 min at 90 V. The RNA was transferred to a positively charged nylon membrane (Roche) using the Biometra Fastblot apparatus at constant current (200 mA) for 30 min. RNA was UV-crosslinked in the Stratalinker 2400 (Stratagene) using the auto-crosslink function 2 times. The membrane was transferred to hybridization tubes and pre-hybridized in 10 mL of ULTRAhyb-Oligo hybridization buffer (Thermo Fisher) at 37°C rotating in a hybridization oven for 1 h. 20 pmol (5 μL of 4 μM stock) of denatured and snap-cooled biotinylated ssDNA oligos complementary to the indicated targets (see Table S1) were added and hybridized overnight rotating at 37°C. The signal was developed using the Chemiluminescent Nucleic Acid Detection Module kit (Thermo Fisher) according to manufacturer’s instructions and recorded on the Amersham Imager 680 (Cytiva).

### RNA extraction and total RNA sequencing

200,000 cells were seeded 24 h before RNA extraction and treated with dTAG13 or DMSO for indicated times. RNA was extracted using Qiagen RNeasy Mini kit (Qiagen) according to manufacturers instructions with on column DNase digestion for 30min using RNase-Free DNase (Qiagen) with 27 Kunitz units per column. Libraries were prepared using the Illumina Stranded Total RNA Library Prep, including a ribosomal RNA depletion step, and sequenced on the Illumina NovaSeq 6000 (51-nt paired-end reads).

### Immunoprecipitation

Immunoprecipitation experiments were performed using ProteinG Dynabeads (Thermo Fisher Scientific). Per sample, 10 µL of beads were washed with 1× PBST and pre-coupled with 1 µL of anti-FLAG M2 antibody (Sigma-Aldrich) for 1 h at room temperature with rotation.

For each replicate, 10 million cells were harvested by trypsinization and washed once with 1× PBS. Cells were lysed in 500 µL of IP buffer (40 mM HEPES pH 7.4, 150 mM NaCl, 2 mM MgCl₂, 5% glycerol, 0.5% Triton X-100) supplemented with 0.5 µL/mL Turbonuclease (Sigma-Aldrich) and 1× Halt protease inhibitor cocktail (Thermo Fisher Scientific). Lysis was performed for 1 h with rotation at 4°C.

Lysates were cleared by centrifugation at 20,000 × g for 30 min at 4°C, and the supernatants were transferred to new tubes. The cleared lysates were incubated with the antibody-coupled beads for 2 h with rotation at 4°C, followed by four washes with IP buffer without Turbonuclease and protease inhibitors. Finally, three washes were performed with buffer containing no detergent or glycerol (40 mM HEPES pH 7.4, 150 mM NaCl, 2 mM MgCl₂), with tube changes after the first wash to minimize detergent contamination.

### Mass spectrometry

Washed beads were resuspended by vortexing in 5 µL of digestion buffer (3 M GuaHCl, 20 mM EPPS at pH 8.5, 10 mM CAA, 5 mM TCEP), and 1 µL of 0.2 mg/mL LysC protease (Promega) in 50 mM HEPES (pH 8.5) was added. Proteins were digested for 2 h at room temperature with rotation. The samples were diluted with 17 µL of 50 mM HEPES (pH 8.5) and digested with 1 µL of 0.2 mg/mL trypsin (Promega) in 0.2 mM HCl at 37°C overnight with interval mixing at 2000 rpm for 30 sec every 15 min.

All samples were acidified with 1.5ul 20% TFA and the supernatant was transferred to injection vials.

#### ADNP co-immunoprecipitation analysis

Mass spectrometry analysis of ADNP co-immunoprecipitation samples was performed using an Orbitrap LUMOS mass spectrometer (Thermo Fisher Scientific) coupled to a VanquishNeo nanoLC system with an Easy Source and a 75 μm × 15 cm EasyC18 column (Thermo Fisher Scientific). Samples were loaded on a C18 (0.3 × 5 mm) trap column and analyzed using backward flush.

Chromatographic separation was performed with an 80-minute gradient as follows: 0-1 min, 2-6% B in A; 1-43 min, 6-20% B; 43-65.5 min, 20-35% B; 65.5-67.5 min, 35-45% B; 67.5-68.5 min, 45-100% B; 68.5-80 min, 100% B. Mobile phase A consisted of 0.1% formic acid in water, and mobile phase B consisted of 0.1% formic acid, 80% acetonitrile in water. The gradient was run at room temperature with a flow rate of 200 µL/min.

#### GTF3C1 co-immunoprecipitation analysis

Mass spectrometry analysis of GTF3C1 co-immunoprecipitation samples was performed using an Orbitrap Fusion LUMOS mass spectrometer (Thermo Fisher Scientific) coupled to an Easy-nLC system with a DPV source and a PharmaFluidics 50 cm column (PharmaFluidics).

Chromatographic separation was performed with a 75-minute gradient as follows: 0-3 min, 2-6% B in A; 3-43 min, 6-22% B; 43-52 min, 22-28% B; 52-60 min, 28-36% B; 60-61 min, 36-80% B; 61-75 min, 80% B. Mobile phase A consisted of 0.1% formic acid in water, and mobile phase B consisted of 0.1% formic acid in acetonitrile. The gradient was run at room temperature with a flow rate of 500 µL/min.

### Purification of TFIIIC from insect cells

A six-gene TFIIIC expression biGBac expression plasmid was obtained from Christoph Müller ^38^. The biGBac expression plasmid was transformed into competent DH10-EmBacY cells and transformed bacteria were recovered in LB medium at 37°C overnight. The recovered bacteria were plated on blue-white selection LB-agar plates containing X-gal, IPTG, kanamycin, tetracycline, and gentamicin. Plates were incubated at 37°C for 24–48 h until blue and white colonies could be clearly distinguished. A white colony was grown in LB medium containing kanamycin, tetracycline, and gentamicin at 37°C for 16 h.

Bacmid DNA was isolated by alkaline lysis using buffers from a Miniprep kit (Qiagen) and isopropanol precipitation. The bacterial pellet was resuspended in 250 μL buffer P1, lysed with 250 μL buffer P2, and neutralized with 350 μL buffer N3. The lysate was cleared by centrifugation at 20,000 × g for 10 min. The supernatant (750 μL) was mixed with 750 μL isopropanol, inverted, and incubated on ice for 5 min. The DNA precipitate was pelleted by centrifugation at 20,000 × g for 20 min at 4°C. The DNA pellet was washed with 70% ethanol and centrifuged at 20,000 × g for 5 min at 4°C.

In a laminar flow hood, the supernatant was removed, the DNA pellet was air-dried and resuspended in 40 μL sterile water. Sf9 insect cells were transfected, the baculovirus was amplified, and protein was expressed using standard procedures as described below.

TFIIIC complex was purified from insect cells as described previously ^38^ with minor modifications. 4 L of insect cells grown in SF-4 Baculo Express ICM (BioConcept) were infected at a density of 0.5 million cells /mL with 20 mL of virus. The cells were grown for 72h after infection or until viability dropped below 80%. After harvesting by centrifugation, the cells were washed once with 1x PBS and flash-frozen in liquid nitrogen before storage at −80°C until use.

Frozen insect cell pellets were resuspended for 2 hours at 4°C while stirring gently in lysis buffer (20 mM HEPES pH 7.5, 500 mM NaCl, 2 mM MgCl₂, 0.1% NP-40, 10% glycerol and 0.25 mM DTT), supplemented with SigmaFast EDTA-free protease inhibitor tablets (Sigma-Aldrich) and 4 μL of benzonase (Sigma-Aldrich) per 50 mL of lysis buffer. For approximately 50 mL of cell pellet, 100 mL of lysis buffer was used.

The resuspended pellet was lysed using a sonicator for 3 minutes. The lysate was then centrifuged at 30,000 × g for 1 hour at 4°C to remove cell debris.

2 mL of Anti-FLAG M2 agarose beads (Sigma-Aldrich) were incubated with the resulting supernatant for 2 hours on a rolling plate at 4°C. Beads were loaded onto an Econo-Column Chromatography column (Bio-Rad) and washed with 60 mL of wash buffer 1 (20 mM HEPES pH 7.5, 500 mM NaCl, 10% glycerol, 0.1% NP-40, and 0.25 mM DTT).

The washed resin was incubated with 2× column volumes (4 mL) of elution buffer 1 (20 mM HEPES pH 7.5, 500 mM NaCl, 10% glycerol, 0.1% NP-40, 0.25 mM DTT, and FLAG peptide (0.2 mg/mL)) for 30 min at 4°C. Wash buffer 2 (20 mM HEPES pH 7.5, 100 mM NaCl, 10% glycerol, 0.1% NP-40, and 0.25 mM DTT) was added to the eluted sample (12 mL) to dilute the initial salt concentration to 200 mM NaCl.

The elution was checked by SDS-PAGE and then applied to a MonoQ 5/50 column (Cytiva) pre-equilibrated in buffer A (20 mM HEPES pH 7.5, 200 mM NaCl, 5% glycerol, and 5 mM DTT). Sample elution was carried out by applying a linear gradient from 0 to 100% buffer B (20 mM HEPES pH 7.5, 1 M NaCl, 5% glycerol, and 5 mM DTT) over 70 mL. Peak fractions were analyzed by SDS-polyacrylamide gel electrophoresis.

The fractions containing TFIIIC were pooled, concentrated up to 2 mg/mL, buffer-exchanged using storage buffer (20 mM HEPES pH 7.5, 200 mM NaCl, and 5 mM DTT), flash-frozen, and stored at −80°C until use.

### Purification of ChAHP from insect cells

Purification of ChAHP was done as in Ostapcuk et al.^34^ in short: Full-length versions of ChAHP subunits were subcloned into pAC8 or pFastBac-derived vectors with and N-terminal His6-tag for ADNP and CHD4 as well as an N-terminal Strep-tag II for HP1γ. Baculoviruses were generated in Sf9 cells using the Bac-to-Bac method for pFastBac-derived vectors or by cotransfection with viral DNA for pAC8-based vectors. 1 L of Hi5 cells coinfected with Baculoviruses encoding for His6-tagged ADNP and Strep-tagged HP1γ was combined with 2 L of Hi5 cells expressing His6-tagged CHD4. Cells were lysed in lysis buffer and the cleared lysate was passed over a 5mL Strep-Tactin Sepharose (IBA) column. The bound complex was eluted in 50 mM Tris-HCl, pH 7.5, 100 mM NaCl, 5 mM β-mercaptoethanol, 2.5 mM desthiobiotin, and bound to an anion-exchange chromatography column (Poros HQ) equilibrated in 50 mM Tris-HCl, pH 7.5, 100 mM NaCl, 5 mM β-mercaptoethanol. The bound proteins were eluted using a linear NaCl gradient, concentrated and further purified by SEC (HiLoad Superdex 200 26/600) in 50 mM HEPES-OH, pH 7.4, 150 mM NaCl and 0.5 mM TCEP. Fractions containing the ChAHP complex were concentrated and reinjected to a Superdex200 10/300 column equilibrated in the same buffer, concentrated again and frozen at −80°C until use.

### In-vitro pulldown of TFIIIC and ChAHP

Recombinant ChAHP and TFIIIC complexes were incubated at 0.75 μM each in a total volume of 50 μL pulldown buffer (20 mM HEPES pH 8.0, 150 mM NaCl, 1.5 mM MgCl₂, 0.05% NP-40) for 15 min on ice. The protein mixture was then incubated with 10 μL Magnetic Strep beads (IBA) for 15 min on ice. The ChAHP complex served as bait due to the StrepII-tag on the HP1γ subunit. Beads were washed three times with pulldown buffer and proteins were eluted by boiling at 75°C for 5 min in 1× SDS loading buffer. Eluted proteins were separated on 4-6% Tris-Acetate gels (Invitrogen) and visualized using InstantBlue protein stain (Abcam). Gels were imaged using a GelDoc system (Bio-Rad). Negative controls included beads alone and individual complexes incubated separately with beads to assess non-specific binding.

### Genomics data analysis

#### SINE curation

Given the short length and proximity of SINEs to each other and other POL III transcription units, we excluded all SINEs which are within 200 bp of another SINE or POL III transcription unit. With an average ChIP fragment size of 200-400 bp, this ensures unambiguous assignment of ChIP peaks to genomic annotations (see Figure S1E).

#### ChIP seq alignment

Reads were mapped to the mouse mm39 genome using STAR version 2.7.3 ^56^ and repeat sensitive settings as suggested by Teissandier et al^57^. The parameters used were STAR: -- outSAMtype BAM SortedByCoordinate --winAnchorMultimapNmax 5000 --alignEndsType EndToEnd --alignIntronMax 1 --alignMatesGapMax 2000 --seedSearchStartLmax 30 -- alignTranscriptsPerReadNmax 100000 --alignWindowsPerReadNmax 100000 --alignTranscriptsPerWindowNmax 300 --seedPerReadNmax 1000 --seedPerWindowNmax 300 -- seedNoneLociPerWindow 1000 --outBAMsortingBinsN 100 --clip3pAdapterSeq CTGTCTCTTATACACATCT AGATGTGTATAAGAGACAG --limitBAMsortRAM 10000000000 -- alignSJDBoverhangMin 999.

#### Peak finding

Peak calling was performed using MACS3 version 3.0.1 ^58^ on combined replicates for each ChIP condition. Read counts within peaks (resized to 300 bp around peak summits) were calculated for both ChIP and Input samples using featureCounts() from the Rsubread package ^59^. Counts were normalized to library size (counts per million, CPM), and peaks were retained if they showed >1.2-fold ChIP/Input enrichment and CPM >1 in all replicates.

#### Coverage plots

ChIP-seq signal tracks were visualized using the MiniChip package (version 0.0.0.9600) in R. Representative genomic regions spanning 1000 bp upstream and downstream of features of interest were selected. Signal tracks were generated from bigWig files using the calcTracks() and plotTracks() functions with 20 bp window smoothing.

#### Heatmaps

ChIP-seq heatmaps were generated using the SummitHeatmap() function from the MiniChip R package (version 0.0.0.9600). Parameters for the individual panels can be found in the code snippets on Github (https://github.com/xxxmichixxx/ADNP_POL3).

#### Metaplots

To generate metaplots we used the CumulativePlots() function of the MiniChip R package (version 0.0.0.9600).

### RNA-seq data analysis

#### Read mapping and quantification

RNA-seq reads were aligned to the mouse genome (GRCm39) using STAR aligner with following parameters: --outFilterType BySJout --outFilterMultimapNmax 100000 –outFilterMismatchNmax --winAnchorMultimapNmax 20000 --alignMatesGapMax 350 --seedSearchStartLmax 30 alignTranscriptsPerReadNmax 30000 --alignWindowsPerReadNmax 30000 -- alignTranscriptsPerWindowNmax 300 --seedPerReadNmax 3000 --seedPerWindowNmax 300 - -seedNoneLociPerWindow 1000--outSAMattributes NH HI NM MD AS nM --outMultimapperOrder Random --outSAMmultNmax 1 --outSAMunmapped Within --limitBAMsortRAM 10000000000. Uniquely mapped reads were quantified at the gene level using featureCounts with the GENCODE vM34 annotation.

For transposable element (TE) analysis, a custom annotation file was created from RepeatMasker data (GRCm39.primary_assembly.repMasker_sensitive.bed). Unlike gene analysis which used only uniquely mapped reads, TE quantification incorporated multimapping reads to account for the repetitive nature of these elements. Read counts were generated using featureCounts with settings allowing for multiple mapping reads, multiple overlaps, and no read fraction assignment. Simple repeats, low complexity regions, and rRNA were excluded from the analysis to focus on biologically relevant transposable elements. Only TEs with at least 120 normalized read counts across samples were included in downstream analyses to focus on robustly expressed elements.

#### Differential expression analysis

Differential expression analysis was performed using DESeq2 for both genes and TEs. For differential expression significance, a fold change threshold of 1.6-fold (log₂FC > 0.7) and adjusted p-value < 0.05 were used. Expression changes were evaluated at each condition compared to the respective controls.

For quantitative analysis of specific repeat families, normalized count data was extracted from the DESeq2 object to allow direct comparison between conditions. Count normalization was performed using DESeq2’s variance stabilizing transformation. Mean normalized counts with standard deviation were calculated for each experimental condition across replicates.

#### MA-plots

RNA-seq data visualization was performed using R (version 4.1.0) with the tidyverse, ggplot2, and specialized biological visualization packages. Differential expression results were visualized using MA plots, with significantly regulated genes and repeat elements highlighted (log₂FC > 0.7 and adjusted p-value < 0.05). For repeat elements, specific families were color-coded by class, with particular emphasis on SINE B2 elements.

For transposable element expression analysis, normalized read counts (DESeq2 normalized) were visualized as bar plots with individual data points representing biological replicates. Error bars indicate standard deviation across replicates.

### Proteomics data analysis

Peptides were identified with MaxQuant version 2.2.0.0 using the search engine Andromeda ^60^. The mouse subset of the UniProt database version 2025_03_12 combined with the contaminant database from MaxQuant was searched, and the protein and peptide false discovery rate (FDR) values were set to 0.05. Statistical analysis was performed using limma within the einProt package (version 0.9.7)^61^. Results were filtered to remove reverse hits, contaminants, and peptides found in only one sample. For visualization and comparison of protein abundances across different conditions, iBAQ (intensity-based absolute quantification) values were used, with log10 transformation applied to normalize the data. Graphs were generated to compare protein abundances across experimental conditions, displaying both mean values and individual replicate data points to illustrate experimental variability. Analyses focused particularly on proteins of the GTF3C complex (GTF3C1-6) and their interactions with ADNP, CBX3, and CHD4 across various experimental conditions.

